# ProteinAligner: A Multi-modal Pretraining Framework for Protein Foundation Models

**DOI:** 10.1101/2024.10.06.616870

**Authors:** Li Zhang, Han Guo, Leah Schaffer, Young Su Ko, Digvijay Singh, Hamid Rahmani, Danielle Grotjahn, Elizabeth Villa, Michael Gilson, Wei Wang, Trey Ideker, Eric Xing, Pengtao Xie

## Abstract

Protein foundation models, particularly protein language models, have demonstrated strong success in learning meaningful representations of proteins using transformer architectures pretrained on large-scale protein datasets with self-supervised learning. These representations have been highly effective for downstream tasks such as predicting protein functions and properties. However, most current protein foundation models focus on pretraining with amino acid sequences, often neglecting additional modalities like protein structures and related literature, both of which provide valuable insights. To address this gap, we propose a multi-modal pretraining approach that integrates three key modalities - protein sequences, structures, and literature text. In our framework, the protein sequence modality serves as the anchor, with the other two modalities aligned to it, enhancing the model’s capacity to capture more comprehensive protein information. ProteinAligner out-performed state-of-the-art protein foundation models in predicting protein functions and properties across diverse down-stream tasks.

## Introduction

Proteins play a fundamental role in virtually all biological processes. Understanding their functions and properties is central to advancing fields such as drug discovery (1), diagnostics (2), and biotechnology (3). Recent advances in artificial intelligence, particularly in transformer-based models (4), have led to the development of protein foundation models capable of learning rich representations from largescale protein datasets (5–11). These models, particularly protein language models (PLMs) (5–7, 10, 11), have shown remarkable success in performing various downstream tasks such as protein function prediction (12, 13), property prediction (14, 15), structure prediction (16, 17), and protein design (18, 19).

Despite these successes, current PLMs predominantly focus on amino acid sequences while overlooking the wealth of complementary information available in other modalities. Protein structures, for example, provide critical three-dimensional information that is essential for understanding how proteins fold and interact with other molecules, directly influencing their biological functions (20). The spatial arrangement of amino acids, which governs interactions such as binding affinities and functional sites, cannot be readily inferred from sequence data alone (21, 22), making the integration of structural data crucial for a more comprehensive understanding of protein behavior. Similarly, the vast amount of biological literature contains experimentally validated insights into protein mechanisms, behavior, and interactions that are often context-specific and difficult to infer from sequences or structures alone (23, 24). Literature captures critical information about post-translational modifications, protein dynamics in various environments, and interaction networks - details accumulated from years of experimental studies. By incorporating these additional modalities - protein structures and related literature - protein foundation models can move beyond sequence prediction to a more robust, context-aware understanding of protein biology. This multi-modal integration has the potential to greatly enhance the representational power of these models, enabling more accurate predictions of protein functions and behaviors in diverse biological scenarios. While some studies have explored the use of two modalities, such as protein sequence and text (24–26) or protein structure and sequence (8, 9, 27), the simultaneous integration of three modalities for pretraining protein foundation models remains underexplored.

To address this gap, we introduce ProteinAligner, a multi-modal pretraining framework that combines protein sequences, structures, and literature text. Our framework aligns these modalities with the protein sequence as the anchor, enabling the model to learn richer and more comprehensive representations of proteins. By integrating diverse protein-related data, ProteinAligner improves the model’s ability to capture intricate biological phenomena, paving the way for more accurate predictions of protein functions and properties. ProteinAligner utilizes three specialized encoders - a sequence encoder, a structure encoder, and a text encoder - to learn representations for each modality. These distinct representations are projected into a shared latent space, enabling direct comparison across modalities. By employing a contrastive alignment strategy (28), ProteinAligner uses protein sequences as the anchor to align corresponding structures and textual descriptions, encouraging similar representations for the same protein and dissimilar representations for different proteins. This approach not only maximizes data utilization by allowing pretraining on incomplete modality data but also captures the comprehensive biological insights provided by each modality. Recently, another independent study (29) also explored learning protein representations across three modalities. Their work and ours were developed independently, with approaches and experiments conceived separately.

ProteinAligner demonstrated superior performance compared to state-of-the-art baselines across various downstream prediction tasks, including detecting type I anti-CRISPR activities, predicting pathogenic missense variants, predicting protein thermostability, identifying potent bioactive peptides, and estimating the minimum inhibitory concentration of antimicrobial peptides.

## Results

### ProteinAligner overview

ProteinAligner is a multi-modal foundation model for protein representation learning, integrating three distinct modalities: amino acid (AA) sequences, 3D structures, and textual descriptions of proteins. ProteinAligner contains three encoders - a protein sequence encoder, a protein structure encoder, and a text encoder - each dedicated to learning representations for its corresponding modality (Fig. 1a). The protein sequence encoder is a protein language model that uses the transformer architecture (4) to extract a representation for the input AA sequence. It represents each AA as a token and employs self-attention (4) to capture long-range dependencies among AAs. The protein structure encoder utilizes Geometric Vector Perceptron Graph Neural Network (GVP-GNN) (30) layers for geometric representation learning of the input protein structure, followed by transformer layers that capture long-range interactions between atomic coordinates. The text encoder employs a transformer architecture, utilizing self-attention to capture long-range dependencies between language tokens. Specifically, we employed ESM (5), a leading protein language model, as the protein sequence encoder, and ESM-IF1 (9) as the protein structure encoder. ESM consists of 33 transformer layers and 650 million parameters, pretrained on 65 million protein sequences. ESM-IF1 features 20 layers and 124 million parameters, pretrained on 12 million computed protein structures and 16,000 experimentally verified structures. The text encoder includes eight transformer layers with a total of 78 million parameters. ProteinAligner uses modality-specific linear projection modules to map the representations extracted by different encoders into a shared latent space with matching dimensions, ensuring that representations from different modalities are directly comparable. ProteinAligner consists of 867 million model parameters in total.

**Fig. 1.**
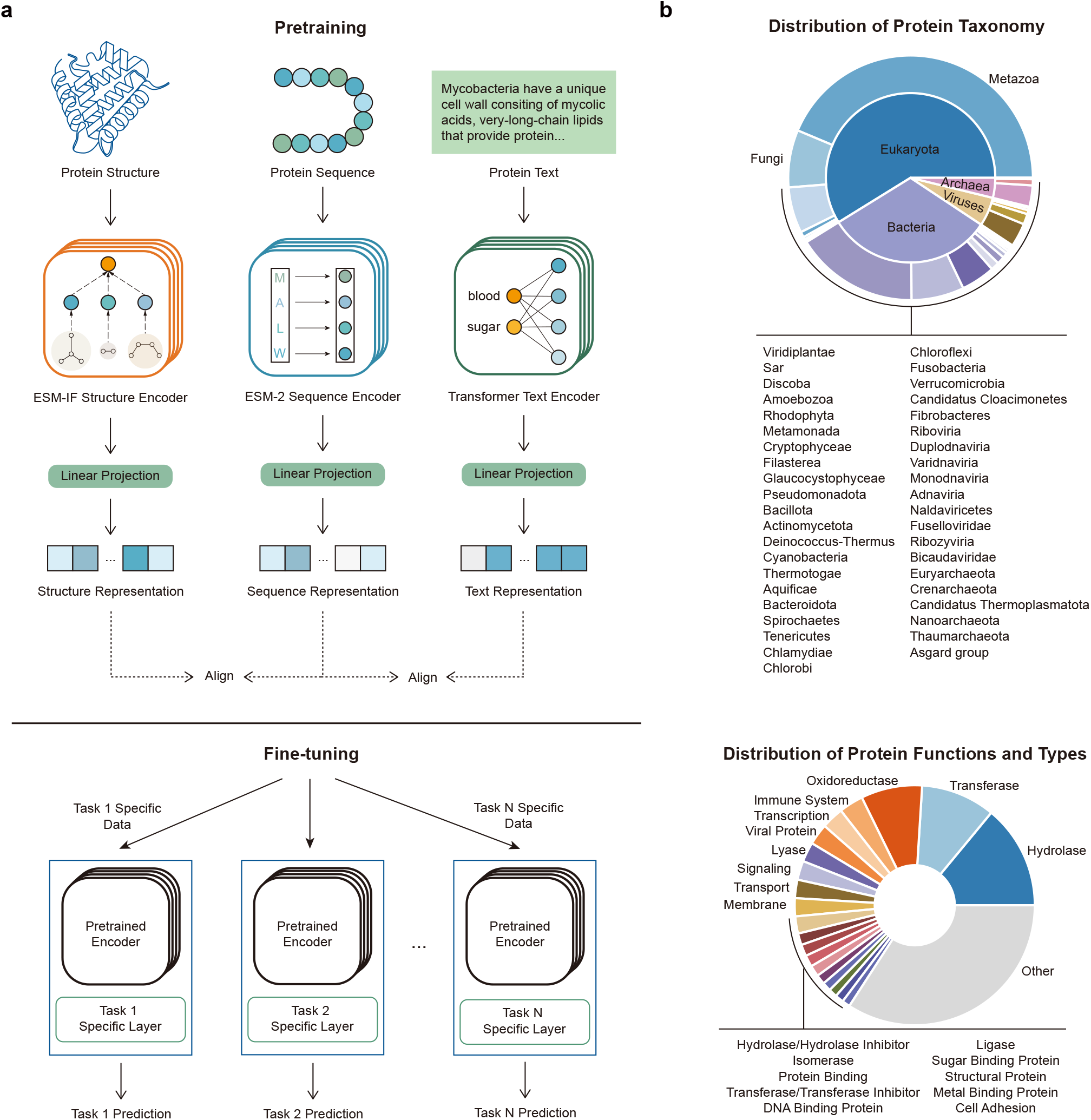
ProteinAligner facilitates multi-modal pretraining of protein foundation models by integrating diverse modalities including amino acid sequences, 3D structures, and textual data. **a**, ProteinAligner consists of three encoders: a protein sequence encoder based on ESM, a protein structure encoder based on ESMIF1, and a transformer-based protein text encoder. These encoders learn representations for protein sequences, structures, and text, respectively. Modality-specific projection modules then transform these representations into a shared latent space, enabling direct comparison across modalities. Using protein sequences as the anchor, ProteinAligner aligns the other two modalities by minimizing contrastive losses. After pretraining, the encoders can be fine-tuned with task-specific data for various downstream applications. **b**, Our curated pretraining data for ProteinAligner spans a diverse range of proteins from various taxonomic groups, functions, and types. The upper chart displays the distribution of protein taxonomy, with the inner ring representing superkingdoms and the outer ring representing kingdoms. The lower chart illustrates the distribution of protein functions and types.

ProteinAligner performs joint pretraining of the three encoders by leveraging a modality alignment strategy, using protein sequences as the anchor modality to align the other two modalities (Fig. 1a). Specifically, given a protein sequence and a protein structure, if they correspond to the same protein, ProteinAligner encourages their representations to be similar, and dissimilar otherwise. The same principle applies for protein text and protein sequences, with representations aligned if they refer to the same protein and separated if not. This alignment is accomplished by minimizing contrastive losses (28, 31) defined on the representations of sequence-structure pairs and sequence-text pairs. ProteinAligner does not require all three modalities to be present simultaneously for each protein in the pretraining data. The alignment can be performed as long as the protein sequence and at least one additional modality - either structure or text - are available. We chose protein sequences as the anchor for alignment because they are the most prevalent data modality in protein databases; nearly every protein has an associated amino acid sequence, whereas information on structures or textual descriptions is often incomplete. By using sequences as the anchor, we can maximize data utilization, ensuring the inclusion of as many proteins as possible in the alignment process. With pretrained encoders in place, they can be fine-tuned on task-specific data to handle a variety of downstream tasks. During this process, the encoders are integrated with task-specific modules, creating models that are customized for specific prediction tasks.

We curated a large-scale pretraining dataset for ProteinAligner by integrating data from the UniProtKB/Swiss-Prot (32) and RCSB PDB (33) databases. The dataset consists of 290,480 proteins, each with an amino acid sequence and a corresponding textual description. 133,726 of them are also associated with protein structures. In total, the dataset contains 133,726 sequence-structure pairs and 290,480 sequence-text pairs. The textual descriptions provide information about the proteins’ functions. Both the structures and the functional descriptions were experimentally validated and reviewed by domain experts. Fig. 1b shows the distribution of protein taxonomy, functions, and types in the dataset.

### ProteinAligner detects type I anti-CRISPR activities

We evaluated the effectiveness of ProteinAligner in detecting type I anti-CRISPR (Acr) activities. Acr proteins are produced by certain viruses, such as bacteriophages, or mobile genetic elements to inhibit the type I CRISPR-Cas immune system in bacteria and archaea (34). The CRISPR-Cas system functions as an adaptive immune mechanism in these microorganisms, recognizing and cleaving foreign DNA from viral invaders. In type I systems, which involve multi-subunit Cas proteins, Acr proteins disrupt this defense by preventing Cas proteins from binding to target DNA or carrying out their cleavage functions. Understanding and detecting these Acr activities is crucial for controlling CRISPR-Cas systems in genetic engineering and leveraging bacteriophages to combat antimicrobial resistance.

Given the amino acid sequences of an Arc protein and a set of Cas proteins from a CRISPR-Cas system, we employed ProteinAligner’s pretrained sequence encoder to extract representation vectors for each protein. These vectors were then input into a convolutional neural network (CNN) based classification module to predict whether the Arc protein could inhibit the CRISPR-Cas system. We utilized the Acr-CRISPR-Cas inhibition dataset (34) for experiments, which comprises 227 pairs of Acr proteins and CRISPR-Cas systems, including 132 experimentally verified positive pairs (Acr inhibits CRISPR-Cas) and 95 negative pairs (Acr does not inhibit CRISPR-Cas). The dataset was randomly split into training and test sets in an 8:2 ratio. We compared ProteinAligner to ESM (5), a protein language model pretrained on protein sequences, and ProtST (24), which builds on ESM by incorporating contrastive pretraining between protein sequences and text. We used area under the ROC curve (AUC), accuracy, precision, recall, and F1 score as evaluation metrics.

ProteinAligner demonstrated significantly better performance across all metrics in detecting type I anti-CRISPR activity compared to ESM (Fig. 2a). For instance, ProteinAligner achieved an AUC of 0.852, substantially surpassing ESM’s AUC of 0.732. Similarly, ProteinAligner’s F1 score of 0.84 was considerably higher than ESM’s F1 score of 0.77. Moreover, ProteinAligner achieved superior performance compared to ProtST (Fig. 2a).

**Fig. 2.**
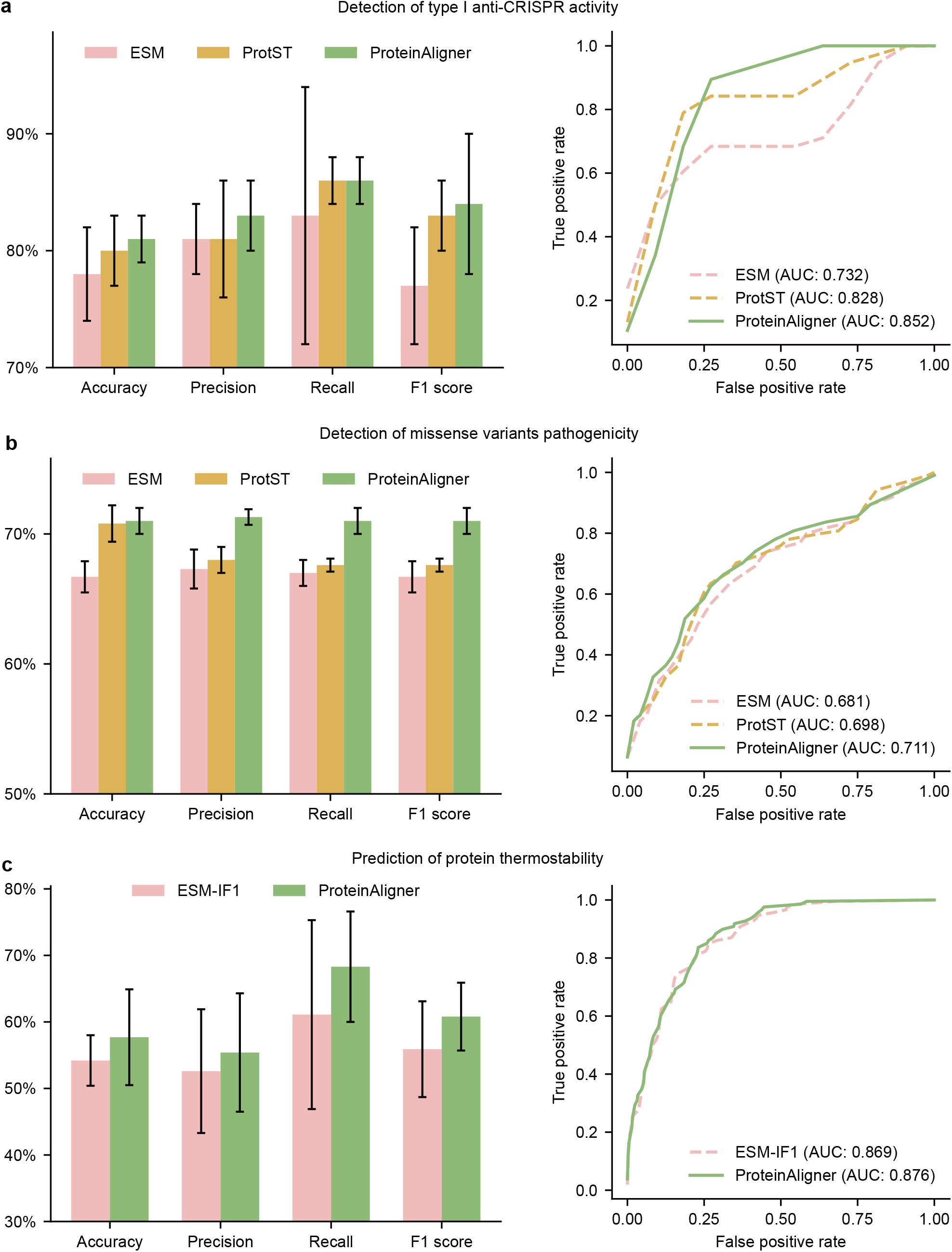
ProteinAligner outperforms state-of-the-art protein foundation models in various downstream tasks. **a-b**, ProteinAligner’s sequence encoder outperforms ESM and ProST in detecting type I anti-CRISPR activity (**a**) and pathogenic missense variants (**b**), achieving higher accuracy, F1 scores (which balance precision and recall), and area under the ROC curve (AUC). **c**, ProteinAligner’s structure encoder outperforms ESM-IF1 in predicting protein thermostability.

### ProteinAligner predicts pathogenic missense variants

Pathogenic missense variants refer to specific types of genetic mutations where a single nucleotide change in a DNA sequence results in the substitution of one amino acid for another in the corresponding protein (35). This change can disrupt the protein’s normal function, potentially leading to diseases or disorders. In the context of pathogenicity, these variants are considered harmful because they alter the protein’s structure or function in a way that impairs biological processes. Depending on the protein’s role, this can lead to a variety of outcomes, from minor effects to severe genetic disorders, such as cystic fibrosis, sickle cell disease, or certain forms of cancer. Identifying and characterizing pathogenic missense variants is crucial in genetic research and clinical diagnostics for understanding inherited diseases and developing targeted treatments.

The inputs for this task are two protein sequences: the wildtype sequence (before mutation) and the mutant sequence (after mutation). We employed the sequence encoder in ProteinAligner to extract representation vectors for both proteins. These vectors were then concatenated and passed through a multi-layer perceptron to predict whether the mutant protein is pathogenic. We used 200 labeled examples from the VariPred (36) dataset, with 100 examples allocated for training and the remaining 100 for testing. ProteinAligner outperformed both ESM and ProtST (Fig. 2b). For example, it achieved an F1 score of 0.71 compared to 0.667 for ESM and 0.676 for ProtST.

### ProteinAligner predicts protein thermostability

Protein thermostability refers to a protein’s ability to maintain its structure and function when exposed to elevated temperatures (37). This characteristic is critical because proteins typically lose their functional shape, or denature, at high temperatures, rendering them ineffective. Thermostability is an important factor in various biological processes and industrial applications. For instance, enzymes with high thermostability are essential in industries such as biotechnology and pharmaceuticals, where reactions often require high temperatures for optimal efficiency. Predicting protein thermostability allows researchers to design or engineer proteins that can withstand challenging conditions, improving their functionality and longevity. Additionally, thermostable proteins are valuable in drug design, as they tend to have better shelf lives and performance under physiological conditions. Accurate predictions of thermostability are crucial for advancing protein engineering and enhancing the reliability of proteins in various applications.

Unlike the previous two tasks, this task takes the 3D structures of proteins, specifically their atomic coordinates, as input. The 3D structure of each protein was processed through ProteinAligner’s structure encoder, generating a representation vector. This vector was then passed through a multi-layer perceptron to predict the protein’s thermostability class. We employed the HP-S^2^C5 dataset (38), which comprises 1,040 proteins spanning five thermostability classes: Hyperthermophilic (above 75°C), Thermophilic (45-75°C), Mesophilic (25-45°C), Psychrophilic (5–25°C), and Cryophilic (*−*20–5°C). 936 proteins were used for training and 104 for testing. We compared ProteinAligner with ESM-IF1 (9), a protein structure encoder pretrained on both protein structures and sequences. ProteinAligner remarkably outperformed ESM-IF1 (Fig. 2c), achieving an F1 score of 0.608 compared to 0.559, and an accuracy of 0.577 compared to 0.542.

### ProteinAligner identifies potent bioactive peptides

Bioactive peptides are short chains of amino acids with specific biological activities (39). They play critical roles in regulating physiological processes, including immune function, metabolism, and cardiovascular health. Identifying bioactive peptides is important because they offer significant potential for developing new therapeutic agents and functional foods. These peptides can serve as natural, targeted treatments with fewer side effects compared to traditional drugs, and their discovery can lead to advancements in both medical applications and nutrition, benefiting public health and disease prevention efforts.

Given the amino acid sequence of a peptide, we employed ProteinAligner’s protein sequence encoder to extract a representation vector, which was subsequently input into a convolutional neural network (CNN)-based classification head to predict whether the peptide has a specific bioactivity. We examined eight distinct bioactivities, including blood-brain barrier penetration (40), umami taste induction (41), antioxidant activity (42), antiviral properties (43), antiparasitic effects (44), T-cell immune response induction (45), inhibition of dipeptidyl peptidase IV (DPP-IV) (46), and modulation of brain activity (47). Given that a peptide can exhibit multiple bioactivities concurrently, we approached each bioactivity prediction as a binary classification task, avoiding the use of a multi-class model that would assign the peptide to a single category. Separate datasets were used for each bioactivity (Methods). The evaluation metrics for this task included accuracy (ACC), balanced accuracy (BACC) (48), sensitivity (SN), specificity (SP), Matthews correlation coefficient (MCC) (49), and the area under the ROC curve (AUC).

ProteinAligner consistently surpassed ESM and ProtST across all eight evaluated tasks (Figs. 3 and 4). For example, in predicting blood-brain barrier penetration, ProteinAligner achieved an AUC of 0.824, outperforming both ESM (0.788) and ProtST (0.767). Similarly, in predicting T-cell immune response induction, ProteinAligner reached an AUC of 0.746, exceeding the performance of ESM (0.678) and ProtST (0.706).

**Fig. 3.**
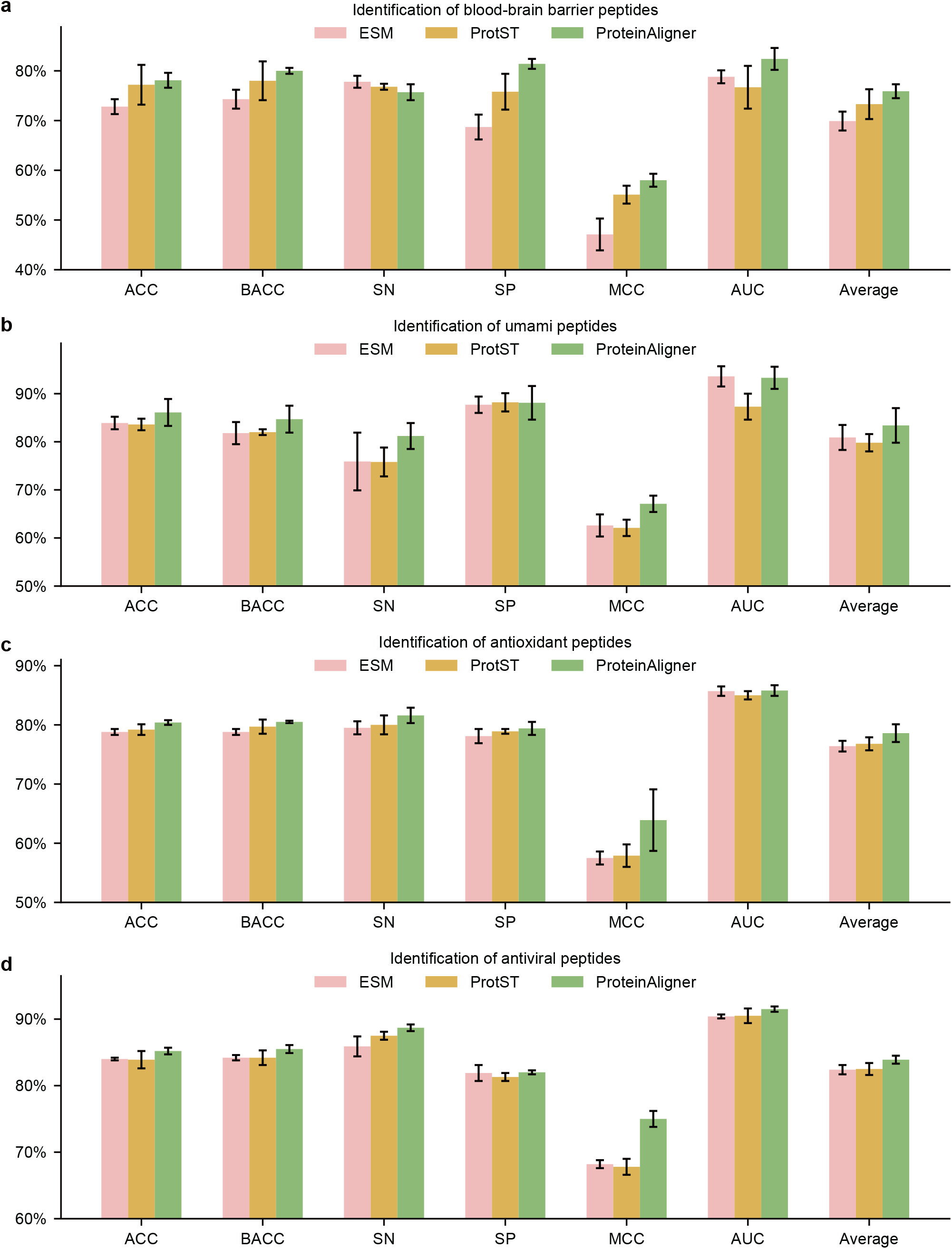
ProteinAligner surpasses ESM and ProtST in identifying potent bioactive peptides. It demonstrated superior performance in predicting blood-brain barrier penetration (**a**), umami taste induction (**b**), antioxidant activity (**c**), and antiviral properties (**d**). Model performance was evaluated using accuracy (ACC), balanced accuracy(BACC), sensitivity (SN), specificity (SP), Matthews correlation coefficient (MCC), and the area under the ROC curve (AUC).

**Fig. 4.**
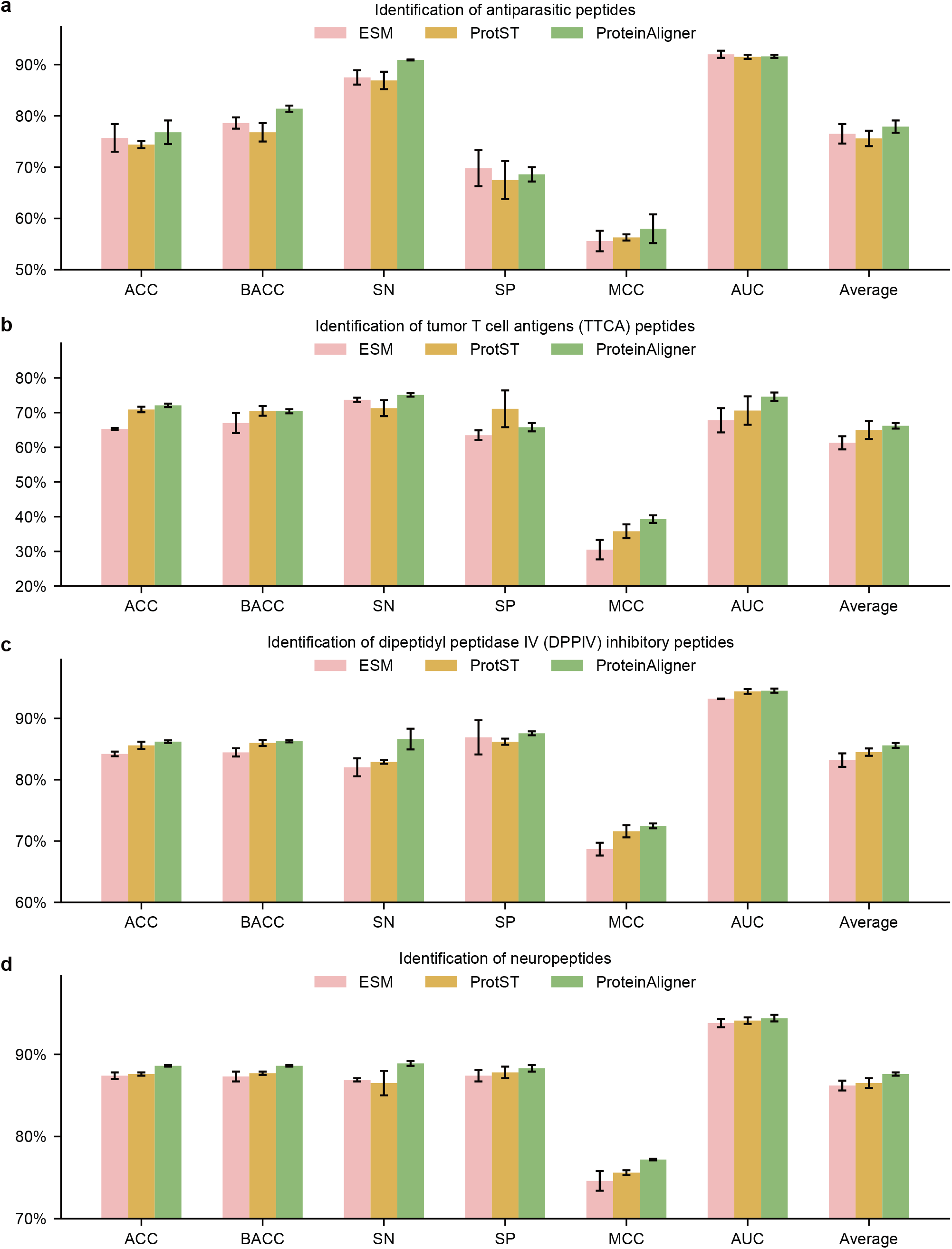
ProteinAligner outperforms ESM and ProtST in identifying potent bioactive peptides. It achieved superior performance in predicting antiparasitic effects (**a**), T-cell immune response induction (**b**), inhibition of dipeptidyl peptidase IV (DPP-IV) (**c**), and modulation of brain activity (**d**). Model performance was assessed using accuracy (ACC), balanced accuracy (BACC), sensitivity (SN), specificity (SP), Matthews correlation coefficient (MCC), and the area under the ROC curve (AUC).

### ProteinAligner predicts the minimum inhibitory concentration (MIC) of antimicrobial peptides

Antimi-crobial peptides (AMPs) are short chains of amino acids that serve as a crucial part of the innate immune response in many organisms, exhibiting broad-spectrum activity against bacteria, viruses, fungi, and even cancer cells (50). They function by disrupting microbial membranes, leading to cell death, and are considered potential alternatives to conventional antibiotics, especially in the face of rising antibiotic resistance. The minimum inhibitory concentration (MIC) is the lowest concentration of an antimicrobial agent, such as an AMP, that prevents visible microbial growth. Accurately predicting the MIC values of AMPs is essential as it allows for the optimization of peptide design for therapeutic use, minimizes potential toxicity, and helps in the early-stage screening of effective peptides before in vitro or in vivo testing. This predictive capability is vital for accelerating the development of AMPs as a novel class of antimicrobial agents in clinical applications.

Given the amino acid sequence of a peptide, we applied ProteinAligner’s protein sequence encoder to extract a representation vector, which was then fed into a multi-layer perceptron-based regression module to predict the MIC of the peptide against a specific pathogen. We focused on Escherichia coli (E. coli), a gram-negative bacterium. We utilized the dataset from (51), comprising 3,695 training and 924 testing examples. Mean squared error was used as the evaluation metric. ProteinAligner achieved lower prediction error compared to ESM and ProtST (Extended Data Fig. 1).

## Discussion

ProteinAligner demonstrates superior performance compared to models like ESM and ProtST by employing a multi-modal approach that integrates sequences, structures, and textual information, offering a more comprehensive understanding of proteins (Figs. 2a, 2b, 3, 4, and Extended Data Fig.1). While ESM focuses on learning from amino acid sequences, ProteinAligner incorporates protein structures, which provide critical insights into folding, stability, and interaction dynamics that sequences alone cannot reveal. Additionally, by drawing from experimental literature, ProteinAligner captures contextual information like functional annotations and post-translational modifications, bridging gaps in sequence and structure-based models. The multi-modal alignment in a shared latent space enables ProteinAligner to generate representations that are not only sequence-based but also grounded in functional and structural knowledge, leading to more accurate predictions. Compared to ProtST, which integrates sequence and text data, ProteinAligner adds the essential dimension of protein structure, crucial for understanding spatial arrangements and physical properties. This enriched representation enables the model to better predict protein functions influenced by structural conformation, outperforming ProtST’s dual-modality approach. Additionally, ProteinAligner surpasses ESM-IF1 (Fig. 2c), which combines sequences and structures, by incorporating textual descriptions that provide further context from experimental literature. ProteinAligner’s contrastive alignment strategy ensures that the representations from sequences, structures, and text are effectively integrated, enhancing its predictive capacity across different tasks. This holistic approach equips ProteinAligner to excel in both sequence- and structure-based predictions, offering a more nuanced understanding of protein properties and delivering superior performance across a range of biological applications.

ProteinAligner’s ability to perform a wide range of prediction tasks presents promising applications across biology, drug discovery, and medicine. In drug discovery, ProteinAligner’s accurate identification of bioactive peptides, such as DPP-IV inhibitors (Fig. 4c), is particularly relevant for developing treatments for metabolic disorders like diabetes. Its capability to predict antimicrobial peptide properties, such as minimum inhibitory concentration (MIC) (Extended Data Fig. 1), is critical for advancing new antimicrobial therapies, especially in addressing the challenge of antibiotic-resistant pathogens. This has important implications for the global fight against antimicrobial resistance. The model’s ability to detect type I anti-CRISPR activities supports the design of more efficient and precise CRISPR-based tools for both research and therapeutic applications (Fig. 2a). Anti-CRISPR systems could be used to enhance the safety of gene editing by mitigating off-target effects or enabling reversible gene modifications. In precision medicine, the prediction of pathogenic missense variants aids in the early detection and diagnosis of genetic disorders. By identifying harmful mutations that may lead to diseases, ProteinAligner can contribute to personalized treatment strategies, improving patient outcomes (Fig. 2b). Additionally, ProteinAligner’s accurate prediction of protein thermostability is vital for protein engineering, biopharmaceutical development, and industrial biotechnology, where stable proteins are necessary for drug formulations and biocatalysts (Fig. 2c). Overall, ProteinAligner’s diverse prediction capabilities position it as a valuable tool that can accelerate innovation in multiple fields, enabling faster therapeutic discoveries, more precise gene-editing tools, and advancements in personalized medicine and protein engineering.

Despite the advantages of ProteinAligner, the model has several limitations. One of the key challenges is the dependency on high-quality structural and textual data, which is not always available for all proteins. While ProteinAligner can perform pretraining even when only sequences and one additional modality (either structure or text) are present, the absence of full multi-modal data for many proteins can limit the model’s ability to learn comprehensive representations. Additionally, ProteinAligner’s reliance on contrastive loss for modality alignment may not fully capture subtle biological nuances in cases where sequence, structure, and text data are not perfectly aligned. Another limitation is the computational cost associated with training multi-modal models, especially when dealing with large-scale protein datasets that involve high-dimensional structural information and large text corpora. Finally, while ProteinAligner improves upon previous models by integrating structure, sequence, and text, it still does not account for other potentially informative modalities, such as protein-protein interactions or functional annotations from various databases, which could further enhance its predictive capabilities.

Future work on ProteinAligner could focus on several key directions to further enhance its performance and applicability. One promising area is the incorporation of additional modalities, such as protein-protein interaction networks and post-translational modifications. These additional data sources could provide deeper insights into protein behavior and interactions, leading to even more robust and comprehensive protein representations. Another direction for future work is to improve the model’s ability to handle incomplete or noisy data by developing more sophisticated alignment strategies that better tolerate inconsistencies between modalities. Enhancing the interpretability of ProteinAligner’s predictions is also a critical area for future research, which could involve incorporating explainability techniques to make the model’s decision-making process more transparent, particularly in cases where sequence, structure, and text data converge. Lastly, expanding ProteinAligner’s applications beyond protein function and property prediction - such as protein design and structure prediction - could broaden its impact across a wide range of biological and biomedical challenges.

**Extended Data Fig. 1.**
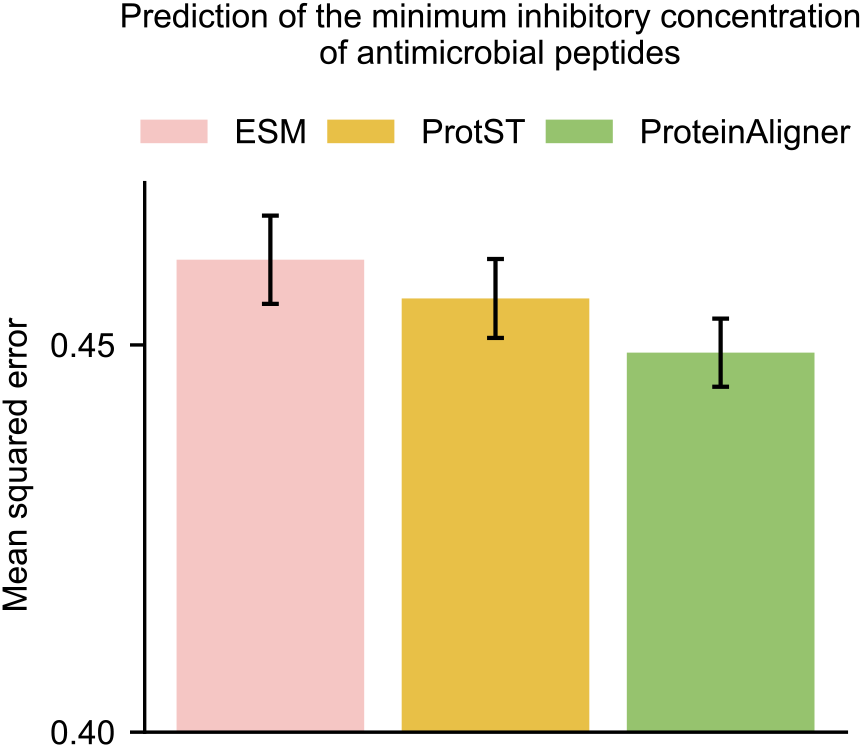
ProteinAligner surpasses ESM and ProtST in predicting the minimum inhibitory concentration (MIC) of antimicrobial peptides, achieving a lower mean squared error.

## Method

### Dataset preprocessing

The pretraining data for ProteinAligner was sourced from the UniProtKB/Swiss-Prot (32) and RCSB PDB (33) databases. UniProtKB/Swiss-Prot is a well-curated repository containing high-quality protein sequences across a wide variety of organisms, along with detailed annotations on protein functions and properties. We utilized version UniProt 2023_02, which was released on May 2, 2023. The RCSB PDB database offers a comprehensive collection of experimentally determined 3D protein structures, derived from methods such as X-ray crystallography, nuclear magnetic resonance (NMR) spectroscopy, and cryo-electron microscopy (cryo-EM). It includes protein structures from a wide range of proteins, such as enzymes, receptors, and antibodies, originating from diverse organisms.

From these databases, we collected sequence-text pairs and sequence-structure pairs (Extended Data Fig. 2). The sequence-text pairs were sourced from UniProtKB/Swiss-Prot. We first obtained a collection of protein entries from Swiss-Prot that included textual descriptions of their functions, by filtering for entries where the `commentType` field was set to ‘Function’. We then retrieved the corresponding sequence for each protein in this collection. Specifically, we accessed the UniProt ID from the `primaryAccession` field and used it to retrieve the corresponding protein FASTA file from the UniProt website, which contains the protein’s sequence. We downloaded all available PDB files from the May 2, 2023 dataset release (33), comprising 200,734 experimentally determined protein structures. We then employed the UniProt ID mapping tool^1^ to link the structures in the PDB files to their corresponding amino acid sequences in the FASTA files. To address memory constraints during pretraining, we excluded protein sequences longer than 300 residues, yielding 133,726 sequence-structure pairs and 290,480 sequence-text pairs.

**Extended Data Fig. 2.**
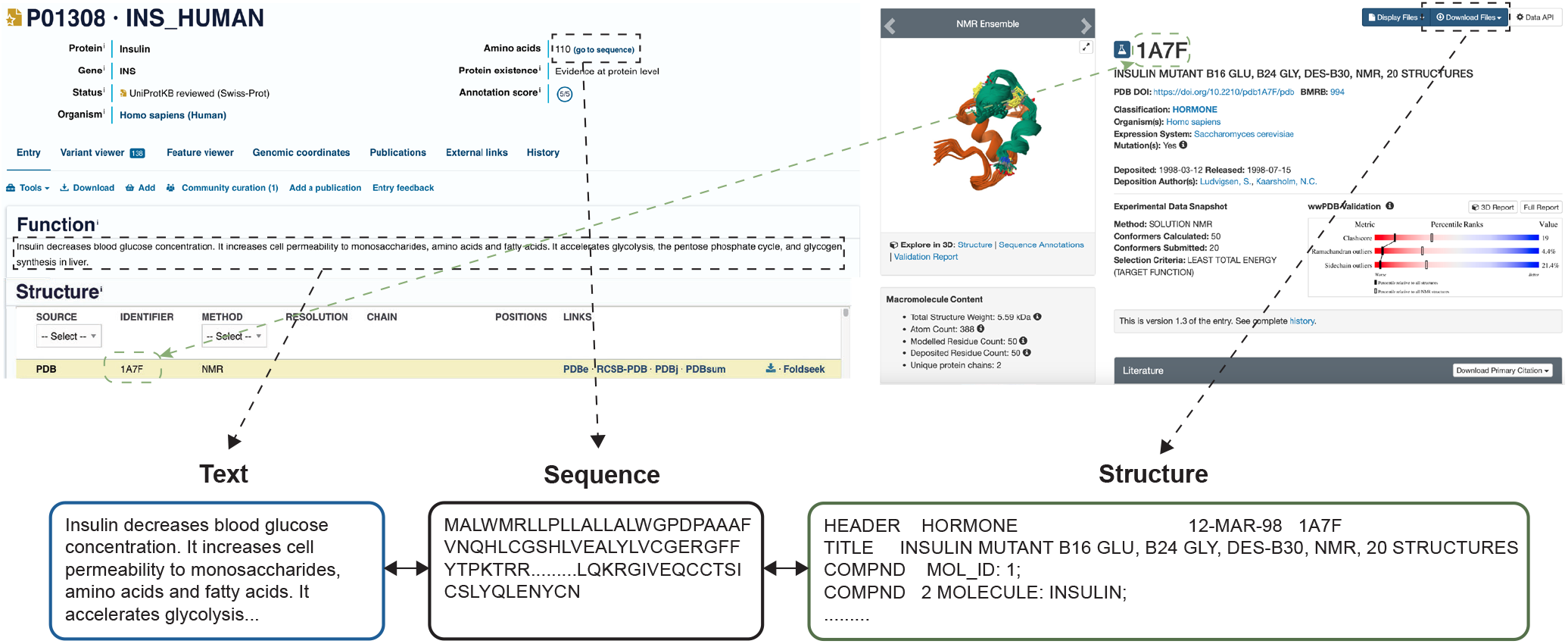
An illustration of constructing sequence-text and sequence-structure pairs from the UniProtKB/Swiss-Prot and RCSB PDB databases. Sequence-text pairs were generated by linking each protein’s amino acid sequence with its functional description, both available on the same UniProtKB/Swiss-Prot webpage. Sequence-structure pairs were created by matching the protein sequences from UniProtKB/Swiss-Prot with their corresponding structures in RCSB PDB, using PDB IDs.

### Encoders in ProteinAligner

ProteinAligner utilizes ESM (16) to learn representations for protein sequences. ESM, a protein language model, was pretrained on 65 million protein sequences from UniRef50 (52) by predicting masked amino acids. The model features 33 transformer layers and an embedding dimension of 1280, allowing it to effectively capture the complexities inherent in protein sequences. To encode protein structures, ProteinAligner employs ESM-IF1 (9), a model trained to address the inverse folding problem - predicting the amino acid sequence from a protein’s backbone atom coordinates. ESM-IF1 comprises an encoder and a decoder, where the encoder extracts a representation vector from the input structure, which is then fed into the decoder to generate the corresponding sequence. ProteinAligner utilizes only the encoder from ESM-IF1, omitting the decoder component. The encoder is composed of four Geometric Vector Perceptron Graph Neural Network (GVP-GNN) (30) layers for geometric feature extraction, followed by eight transformer encoder layers to capture long-range interactions between these features. ESM-IF1 was trained on 12 million AlphaFold2 (8) computed protein structures and 16,000 experimentally verified structures, along with their associated sequences from the UniRef50 dataset (52). The text encoder is a transformer model comprising eight layers and a total of 78 million parameters.

### ProteinAligner pretraining

Given a protein sequence *S* and a protein structure *R*, we employ the sequence encoder *E*_*s*_(*·*) and structure encoder *E*_*r*_(*·*) to extract representation vectors **s** = *E*_*s*_(*S*) and **r** = *E*_*r*_(*R*) for *S* and *R*, respectively. To ensure that the representations are similar when *S* and *R* belong to the same protein, and dissimilar when they do not, we minimize the InfoNCE (31) contrastive learning loss:

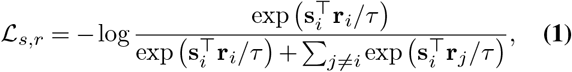

where **s**_*i*_ and **r**_*i*_ represent the sequence and structure representations of the same protein *i*, while **s**_*i*_ and **r**_*j*_ represent the representations of different proteins *i* and *j*. This loss function encourages the alignment of **s**_*i*_ and **r**_*i*_ and discourages the similarity between **s**_*i*_ and **r**_*j*_. The temperature parameter *τ* controls the sharpness of the softmax distribution.

Similarly, given a protein sequence *S* and a protein text description *T*, we use the sequence encoder *E*_*s*_(*·*) and the text encoder *E*_*t*_(*·*) to obtain representation vectors **s** = *E*_*s*_(*S*) and **t** = *E*_*t*_(*T*) for *S* and *T*, respectively. To ensure that the representations are similar when *S* and *T* correspond to the same protein, and dissimilar when they do not, we minimize the following InfoNCE contrastive learning loss:

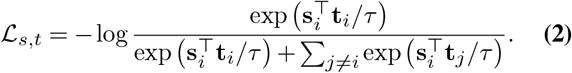

During pretraining, we minimized the sum of the two loss functions with equal weights. The temperature parameter *τ* was configured to 0.07. Pretraining was carried out over 20 epochs with total batch size of 80 using 40 A100 GPUs. We optimized the model weights using the AdamW optimizer (53), with an initial learning rate of 5 *×* 10^−6^, weight decay of 1 *×* 10^−4^, and betas of (0.9, 0.95). The learning rate was dynamically adjusted throughout pretraining via the CosineAnnealingLR scheduler (54).

### Type I anti-CRISPR activity detection

The overall model architecture for this task is illustrated in Extended Data Fig. 3a. The sequence encoder pretrained by ProteinAligner was fine-tuned using data specific to this task. The classification module was based on a CNN which was composed of two 1D convolutional layers, each with a stride of 1 and a kernel size of 7. The first convolutional layer takes an input of 1280 channels and outputs 4 channels. The second convolutional layer maintains the same input and output dimensions. Batch normalization (55) and ReLU activation (56) are applied after each convolutional layer. Following the convolutional layers, two fully connected layers, each with a hidden size of 4, are employed for final classification.

During training, we optimized model weights using the Adam (57) optimizer with an initial learning rate of 3 *×* 10^*−*3^, over a maximum of 250 epochs with a batch size of 32. To prevent overfitting, we employed early stopping when the decrease in training loss fell below 0.005 and applied weight decay, starting at 0.01 and gradually reducing to 0.001. We also performed a hyperparameter sweep on the dropout rate (58), exploring values between 0.3 and 0.5. Additionally, we implemented a learning rate reduction strategy, decreasing the rate by a factor of 0.9 if validation performance did not improve for 10 consecutive epochs.

The metrics used to evaluate model performance in this task include accuracy, precision, recall rate, and F1 score. These metrics are calculated based on the number of true positives (TP), false positives (FP), false negatives (FN), and true negatives (TN), and are defined as follows:

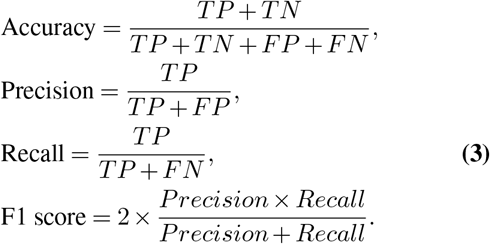

### Pathogenic missense variants prediction

The overall model architecture is depicted in Extended Data Fig. 3b. For this task, the sequence encoder pretrained by ProteinAligner was fine-tuned. The classification module was based on a multi-layer perceptron, which comprises a fully connected layer with a hidden state size of 1280, a dropout layer with a probability of 0.5, a leaky ReLU activation (59), and a second fully connected layer with a softmax activation function. We employed the Adam optimizer with a learning rate of 1 *×* 10^−4^and a weight decay of 1 *×* 10^−3^, training the model for a maximum of 200 epochs with a batch size of 32. To mitigate overfitting, we applied an early stopping strategy: if validation performance did not improve over 10 consecutive epochs, training was halted, and the model checkpoint with the best validation performance was retained as the final model. The model’s performance was evaluated using accuracy, precision, recall, and F1 score, as defined in Equation 3.

### Thermostability prediction

The overall model architecture is depicted in Extended Data Fig. 3c. The structure encoder, pretrained using ProteinAligner, was fine-tuned for this task. The classification module, implemented as a multilayer perceptron (MLP), includes a fully connected layer with a hidden dimension of 128, followed by layer normalization (60), a ReLU activation, and a second fully connected layer to produce the classification logits. For model optimization, we employed the AdamW (53) optimizer with a learning rate of 2 *×* 10^*−*2^, a batch size of 4, and a weight decay of 5 *×* 10^*−*2^. A one-cycle learning rate scheduler (61) was applied, and the model was trained for 8 epochs. Performance was evaluated based on accuracy, precision, recall, and F1 score, as defined in Equation 3.

### Identification of potent bioactive peptides

The overall model architecture is illustrated in Extended Data Fig. 3d. The sequence encoder, pretrained by ProteinAligner, was fine-tuned for each of the eight tasks. The classification module employs a convolutional neural network (CNN) with six layers, structured as follows: a 1D convolutional layer (kernel size = 3, stride = 1, padding = 2), followed by BatchNorm and ReLU activation; a max pooling layer (kernel size = 2, padding = 1) and a dropout layer with a probability of 0.15; another 1D convolutional layer (kernel size = 3, stride = 1, padding = 2), followed by BatchNorm and ReLU activation; a max pooling layer (kernel size = 2, padding = 1) and a dropout layer with a probability of 0.15; a fully connected layer with a hidden state size of 64, followed by ReLU activation and a dropout layer (probability = 0.15); and finally, a fully connected binary classification layer with sigmoid activation.

**Extended Data Fig. 3.**
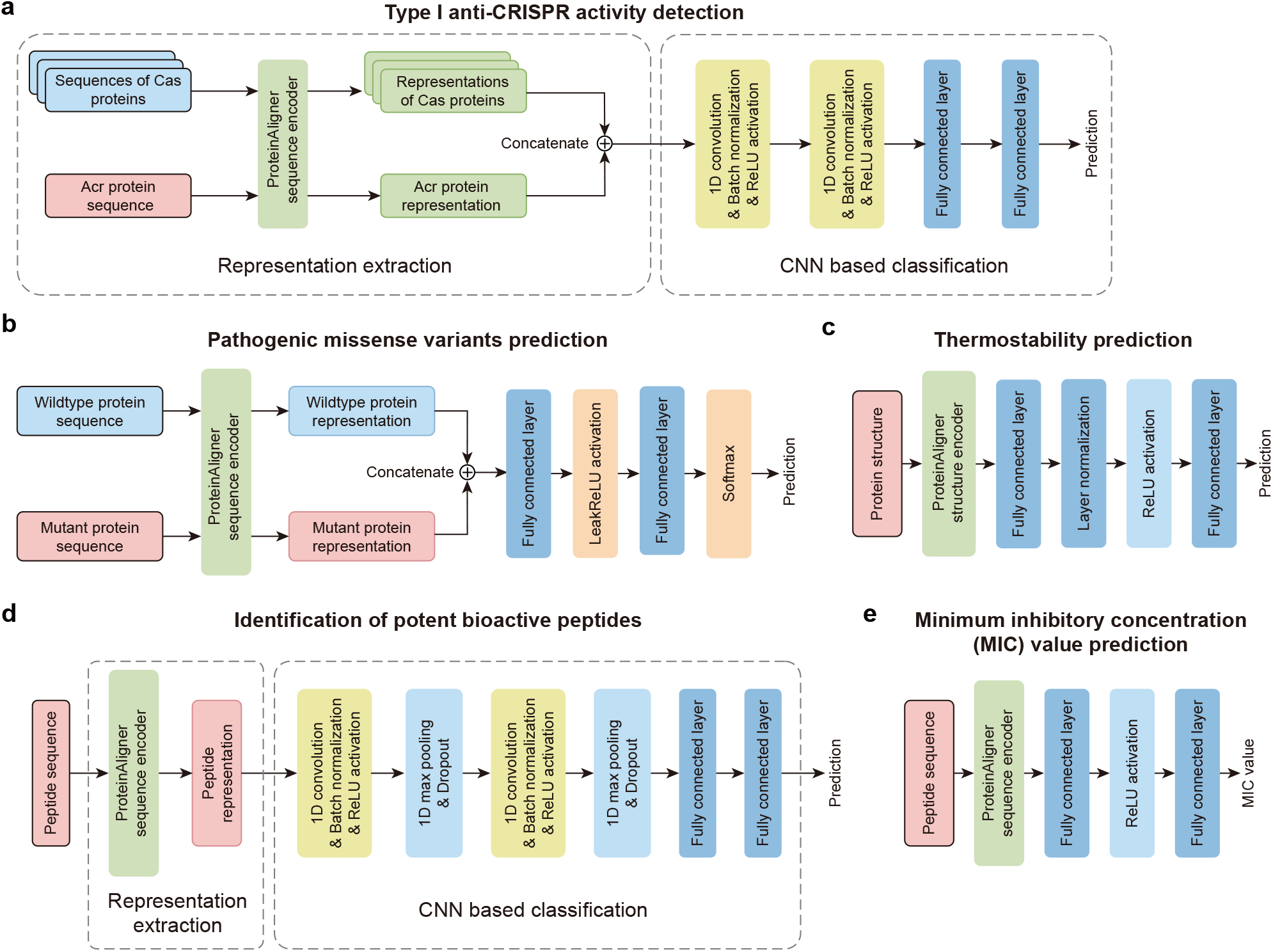
Model architectures used in downstream tasks. **a**, Model architecture used in type I anti-CRISPR activity detection. **b**, Model architecture used in pathogenic missense variants prediction. **c**, Model architecture used in thermostability prediction. **d**, Model architecture used for identifying potent bioactive peptides. **e**, Model architecture used for predicting minimum inhibitory concentration values.

In the blood–brain barrier peptide (BBP) prediction task, the objective is to classify whether a peptide can penetrate the blood–brain barrier (i.e., BBP) (40). We employed the BBPpred dataset (40), consisting of 100 BBPs and 100 non-BBPs for training, and 19 BBPs and 19 non-BBPs for testing. In the umami peptide prediction task, the goal is to determine whether a peptide elicits an umami taste (41). For this task, we used the iUmami-SCM dataset (62), with a training set of 112 umami peptides and 241 non-umami peptides, and a test set of 28 umami peptides and 61 non-umami peptides. In the antioxidant peptide prediction task, the aim is to classify peptides based on their antioxidant properties (42). We used the AnOxPePred dataset (63), containing 582 antioxidative peptides and 241 non-antioxidative peptides for training, with a test set comprising 28 antioxidative peptides and 61 non-antioxidative peptides. For the antiviral peptide prediction task, the objective is to predict whether a peptide has antiviral activity (preventative and therapeutic against viral infections) (43). We utilized the ABPDiscover dataset (64), which includes 2321 antiviral peptides and 2321 non-antiviral peptides for training, and 623 antiviral peptides and 623 non-antiviral peptides for testing. In the antiparasitic peptide prediction task, the goal is to identify peptides with antiparasitic activity (44). Using the PredAPP dataset (65), we trained on 255 antiparasitic peptides and 255 non-antiparasitic peptides, and tested on 46 antiparasitic peptides and 46 non-antiparasitic peptides. The tumor T cell antigen prediction task aims to classify peptides capable of inducing a T-cell immune response (45). We used the iTTCA-Hybrid dataset (45), including 470 antigenic peptides and 318 non-antigenic peptides for training, with 122 antigenic peptides and 75 non-antigenic peptides for testing. For the dipeptidyl peptidase IV (DPP-IV) inhibitory peptide prediction task, the goal is to identify peptides that inhibit DPP-IV activity (66). We used the iDPPIV-SCM dataset (66), containing 532 inhibitory peptides and 532 non-inhibitory peptides for training, and 133 inhibitory peptides and 133 non-inhibitory peptides for testing. Lastly, in the neuropeptide (NP) prediction task, the aim is to classify peptides as neuropeptides or non-neuropeptides (47). We used the PredNeuroP dataset (47), containing 1940 neuropeptides and 1940 non-neuropeptides for training, and 485 neuropeptides and 485 non-neuropeptides for testing.

During training, we optimized the model using stochastic gradient descent (SGD) with a learning rate of 1 *×* 10^*−*2^, momentum of 0.5, and no weight decay, over 200 epochs. Additionally, we applied step decay for learning rate adjustment and utilized early stopping based on validation accuracy, halting training if no improvement was observed for 40 consecutive epochs. Model performance was assessed using several metrics, including accuracy (ACC), balanced accuracy (BACC) (48), sensitivity (SN), specificity (SP), Matthews correlation coefficient (MCC) (49), and area under the ROC curve (AUC). These metrics were derived from the counts of true positives (TP), false positives (FP), false negatives (FN), and true negatives (TN), and are defined by the following equations:

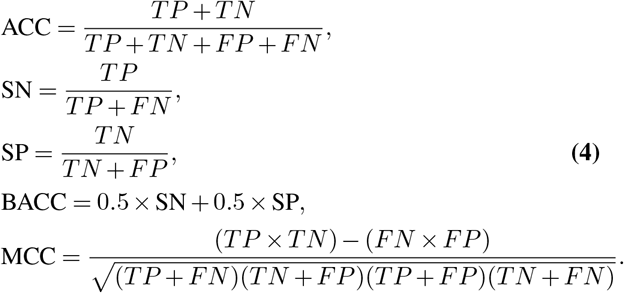

### Minimum inhibitory concentration (MIC) value prediction

The model architecture is illustrated in Extended Data Fig. 3e. The sequence encoder, pretrained with ProteinAligner, was fine-tuned to address this task. The classification module is a multi-layer perceptron (MLP) consisting of two fully connected layers, with a hidden size of 256 and a ReLU activation function. We employed the Adam optimizer with an initial learning rate of 1 *×* 10^*−*4^, training the model for 200 epochs. Throughout the training process, the learning rate was dynamically adjusted at each epoch using the LambdaLR (67) scheduler. To assess the model’s performance, we used mean squared error (MSE) as the evaluation metric, defined as:

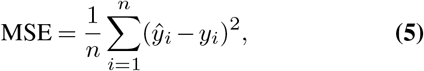

where *n* denotes the number of examples, 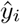 represents the model’s prediction for the *i*-th example, and *y*_*i*_ is the corresponding ground-truth value.

## Data availability

The FASTA and PDB datasets are publicly available in UniProtKB Swiss-Prot and RCSB PDB, respectively. The FASTA and PDB entries for protein sequences and structures in the pretraining data, along with their textual descriptions, are available at https://drive.google.com/file/d/1Ff28ajdyUDQl9JtSNQcUz-6Nbz7FXdEA/view?usp=sharing All the data used in downstream tasks is also publicly available. The data used in type I anti-CRISPR activity detection is available at AcrTransAct. The data for predicting the pathogenicity of missense variants is available at VariPred. The data used in thermostability prediction is available at HotProtein. The data used in peptide bioactivity prediction is available at UniDL4BioPep. The data for predicting the minimum inhibitory concentration (MIC) of antimicrobial peptides is available at DeepAMP.

## Code availability

The source code for ProteinAligner pretraining, along with the pretrained checkpoints, is available at https://github.com/Alexiland/ProteinAligner Additionally, links to the code for downstream tasks can be found in the README file.

## Footnotes

1 https://www.uniprot.org/id-mapping

